# Benchmark validity in graph neural network scoring of metabolic reaction activity on Recon3D: detecting label leakage, memorized noise and input-invariant models

**DOI:** 10.64898/2026.09.02.748315

**Authors:** Thiptanawat Phongwattana, Jonathan H. Chan

## Abstract

Context-specific genome-scale metabolic modeling begins with scoring which of the ~10,600 human reactions are active in a patient’s tumor. Methods in this literature are routinely benchmarked against activity labels obtained by thresholding the same transcriptomic matrix that is supplied to the model as input. We report a self-audit of our own graph attention scorer, MetaGNN, evaluated on TCGA colorectal (*n* = 624), breast (*n* = 1,095) and lung adenocarcinoma (*n* = 517) cohorts, in which two independent failure modes produced a near-ceiling benchmark score and a positive architectural result, neither of which survived inspection. First, under expression-thresholded supervision the framework reaches AUROC 0.9864 ± 0.0008 on TCGA-BRCA. That figure partitions into 5,925 reactions whose labels are a deterministic threshold of the model’s own input, where ranking by the cohort-mean input alone gives AUROC 1.000; and 4,675 reactions whose stored labels we reproduce bit for bit from a seeded pseudo-random number generator, where the model nonetheless reaches 0.9291 ± 0.0030 by memorizing a patient-invariant label vector that patient-level splitting leaves fully visible during training. Second, on the cohort supervised independently of the input, the archived models never received patient data at all. Their released feature tensors are uniformly zero, and independently trained models show no agreement on which patient deviates where (|*r*| ≤ 0.004 on per-patient output residuals, against *r* = +0.32 between output and input residuals on expression-bearing reactions for a model with verified features). A dispersion ratio comparing between-patient output spread against Monte Carlo Dropout sampling spread sits at 1.02–1.03 for all three configurations, against a no-signal null of 1.02 and 2.44 for the verified model. We therefore withdraw a +0.105 AUROC gain attributed to relational edges in an earlier draft of this work. Retraining on rebuilt, verified features gives AUROC 0.5800 ± 0.0017, below both the raw-expression baseline of 0.6342 ± 0.0058 that we establish for this cohort and an information-free indicator baseline of 0.6085. Zero-shot transfer of the BRCA model is at or below chance on METABRIC microarray (0.4926 ± 0.0113, *n* = 200) and on same-platform CPTAC-BRCA RNA-seq (0.4986, *n* = 106). We release the code, the curated colorectal cohort, a script that replays the label vector from its generating seed, and the screening checks we now run before reporting any score. Source code: https://github.com/thiptanawat/MetaGNN-Framework (MIT).

**Highlights:**

- **A near-ceiling benchmark score with no biological content**. Under cohort-shared, expression-thresholded supervision the framework reaches AUROC 0.9864 ± 0.0008 on TCGA-BRCA. It partitions into 1.0000 on the 5,925 reactions whose label is a deterministic threshold of the model’s own input, and 0.9291 ± 0.0030 on the 4,675 reactions whose labels are pseudo-random.
- **The randomized partition is the airtight part of the audit**. We reproduce 4,675 of the 10,600 stored BRCA labels bit for bit from a seeded pseudo-random number generator, and we release the script that does it. A model scoring 0.93 against labels that encode nothing is memorizing a vector that patient-level splitting never hides. Any benchmark whose labels are shared across patients admits this failure mode, whatever the labels contain.
- **A model that never read its input, and three checks that reveal it**. The released colorectal feature tensors are uniformly zero. Independently trained models agree on nothing patient-specific (|*r*| ≤ 0.004 between their per-patient output residuals), and a dispersion ratio of between-patient output spread to MC-Dropout sampling spread sits at 1.02–1.03 for all three archived configurations, against a no-signal null of 1.02 and 2.44 for a model with verified features.
- **We withdraw our own positive result**. The +0.105 AUROC gain we had attributed to reaction–reaction relational edges came from a run that changed the edge topology, the input feature set and the parameter count together. That feature set was never archived, and it shows no patient signal. It is not evidence about network structure.
- **Nothing beat the input, and the input barely beat nothing**. On the cohort with input-independent labels, raw expression alone attains AUROC 0.6342 ± 0.0058, of which an information-free indicator of which reactions carry any expression already supplies 0.6085. The one configuration we retrained on verified features reaches 0.5800 ± 0.0017, below both.
- **What we now run before reporting a score**. Compare against the raw input feature and against an information-free partition indicator; report the median inter-patient output correlation; check that the output depends on the input at all. Each costs minutes. We release all of them with the curated 624-patient cohort and its rebuilt gene-identifier mapping (94.9 % of GPR-bearing reactions resolved).

## 1 Introduction

Context-specific genome-scale metabolic modeling begins with a scoring step: deciding, for a given patient, which of the roughly 10,600 reactions in the human reference network are active. Every downstream analysis, flux balance calculation, biomarker search and vulnerability map inherits whatever that step produces. Production tools take varied approaches to it. GIMME [1] and iMAT [2] score reactions from the expression of their own encoding genes; INIT [3] uses protein abundance as its primary evidence; FASTCORE [4] takes a user-supplied core set and grows it by a network-level optimization; CORDA [5] assigns reactions to confidence tiers from expression and then adds dependencies required to support them; mCADRE [6] ranks candidate reactions by expression evidence together with weighted connectivity to adjacent reactions. Whether and how network context should inform the score is already contested.

It is also hard to settle empirically. Machado and Herrgård [7] found that no transcriptomics-integration method outperformed the others across conditions, and that parsimonious flux balance analysis using no expression data at all was often as good or better when scored against ^13^C flux measurements. Opdam et al. [8] found that the choice of extraction method had the greatest influence on gene-essentiality prediction accuracy, while the expression threshold had the greatest influence on which reactions entered a model at all; they also observe that some algorithms eliminate reactions carrying highly expressed genes when those reactions sit in pathways whose other members are lowly expressed. Gopalakrishnan et al. [9] report that extraction method and expression threshold are the dominant influences on model size, and that the scope of alternate solutions depends on the extraction method and on the topology of the parent model.

### 1.1 The evaluation problem

Underneath that disagreement lies a measurement problem, and it is the subject of this paper. Reaction activity is not observed directly at scale in patient tissue. In its absence, the field’s standard practice is to construct activity labels by thresholding a transcriptomic matrix, and then to evaluate a scorer that was given the same matrix as input. When the label is a deterministic function of the input, a sufficiently flexible model can score near the ceiling by reconstructing the threshold, and that number will be reported as accuracy.

We set out to build a graph-based scorer and to test it against that convention. In evaluating it we found not one such artifact but three, each invisible to the checks we had in place, and each detectable by a diagnostic costing minutes. Rather than report the near-ceiling figures our pipeline produced, we report what they turned out to be.

### 1.2 What this paper does

MetaGNN is an open-source heterogeneous graph attention framework that propagates patient transcriptomes through the Recon3D bipartite graph via GATv2 layers [10] and emits per-reaction scores with Monte Carlo Dropout dispersion [11]. We use it here as an instrument, and report four things.

1. **Under expression-derived supervision, a near-ceiling score with no biological content**. The framework reaches AUROC 0.9864 ± 0.0008 on TCGA-BRCA and 0.9839 ± 0.0005 on TCGA-LUAD. Section 3.2 partitions the BRCA figure: on the 5,925 reactions receiving expression, the label is a threshold of the input and ranking by the input alone gives 1.000; on the remaining 4,675, whose stored labels we reproduce bit-exactly from a seeded generator, the model still reaches 0.9291 ± 0.0030 by memorization. Per-reaction outputs are near patient-invariant, at median inter-patient correlation 0.995.
2. **Two checks for whether a model uses its input at all**. Section 3.3 reports whether independently trained models agree on which patient deviates where, and a dispersion ratio comparing between-patient output spread against the model’s own Monte Carlo Dropout sampling spread. Under the estimators our pipeline uses that ratio has a no-signal null near 1.02; our three archived CRC configurations sit at 1.023, 1.024 and 1.033, while a model with verified per-patient features sits at 2.44. The released feature tensors for that cohort are uniformly zero, and we withdraw the configuration comparison and the relational-edge gain we had drawn from them.
3. **Under supervision independent of the input, nothing beat the input**. On 624 TCGA colorectal patients labeled from Human Metabolic Atlas reconstructions [12], raw expression alone attains AUROC 0.6342 ± 0.0058, and an information-free indicator of which reactions carry any expression already attains 0.6085. Retraining on rebuilt, verified features reaches 0.5800 ± 0.0017, below both.
4. **Transfer fails across consortia, not merely across platforms**. Zero-shot AUROC is 0.4926 on METABRIC microarray and 0.4986 on CPTAC-BRCA, which is the same assay platform as the training data. Four discriminators weigh against platform distribution and label construction as the dominant mechanism, and constrain without excluding a per-gene batch mechanism. Orthogonal biological checks are also weak: subsystem-level concordance with literature expectations is 3 of 8 on BRCA and 0 of 5 on LUAD.

We draw a methodological conclusion rather than a performance claim. Reported gains in this literature should be measured against an explicit raw-expression baseline, on labels that are not a function of the model input and whose provenance has been audited, with an input-dependence check on the trained model and cross-consortium evaluation before any generalization claim. We release the code, the curated colorectal cohort and all of these checks so that the comparison can be applied to other methods, including by readers who wish to apply it to ours. METABRIC and CPTAC are not redistributable, so for those we release preprocessing wrappers rather than data.

### 1.3 Scope and what we do not claim

We do not claim that MetaGNN is a validated patient-specific scoring primitive; on the present evidence it is not. We do not claim that relational edges help, and we explicitly withdraw the earlier version of that claim. We do not compare our AUROC against published reconstruction accuracies, which are computed on different targets and are not commensurate with a reaction-ranking AUROC. No cohort in this study is evaluated against measured reaction fluxes. Nor do we claim that the specific defects we found are present in other groups’ pipelines: two of the three are our own implementation errors. What generalizes is not the errors but the condition that hid them, which is that none of our checks examined the relationship between a model and its own input, and the observation that the checks which would have exposed them are cheap.

## 2 Methods

### 2.1 Module 1: Cohorts and Label Construction

#### Reference network

Recon3D v3 [13] was retrieved from the Virtual Metabolic Human repository in SBML format. Parsed with COBRApy following the constraint-based reconstruction conventions of Thiele and Palsson [14], the model used throughout this work contains 10,600 reactions, 5,835 metabolites and 2,248 genes, which are the counts already present in the distributed SBML file; our flux-consistency check removed nothing. Of the 10,600 reactions, 5,938 carry a gene–protein–reaction (GPR) rule and 4,662 do not. The number of reactions that actually receive a nonzero expression feature is smaller still and depends on identifier-mapping coverage: 5,925 on TCGA-BRCA and 5,635 on the rebuilt TCGA-CRC features. Throughout we partition by the empirical criterion, nonzero expression in at least one patient, rather than by the nominal GPR flag, because it is the partition the model actually experiences.

#### TCGA colorectal cohort (*n* = 624)

TCGA-COAD/READ primary tumors were retrieved from the GDC Data Portal. The download log records 701 files returned by the API and 647 retained after the primary-tumor filter, yielding 624 unique patients after deduplication. Microsatellite instability status was obtained from cBioPortal with complete coverage (487 MSS, 92 MSI-H, 44 MSI-L, 1 not evaluable). Expression is log_2_(TPM + 1) over 21,263 genes. (The stored HDF5 dataset carries the legacy name vst_expression; the transform applied is TPM, not a variance-stabilizing transformation, and we state it here because the raw-expression baseline of Section 3.1 is computed on it.)

#### TCGA-BRCA (*n* = 1,095) and TCGA-LUAD (*n* = 517)

STAR gene counts and clinical tables were retrieved from the GDC Data Portal for both cohorts and processed through the identical pipeline.

#### External cohorts

METABRIC [15] comprises 1,992 tumors (997 discovery and 995 validation) profiled on Illumina HT-12 v3 microarrays; 200 patients were sampled to bound compute cost. CPTAC-BRCA supplies an RNA-seq cohort of 106 patients on the same assay platform as TCGA, which is what makes it the informative control in Section 3.5. For both external cohorts the evaluation labels are constructed by applying the BRCA labeling rule to the external expression matrix; Section 3.5 also reports transfer scored against the training-time label vector, which separates label construction from representation transfer.

#### Two supervision regimes, and why the distinction is the point

The cohorts do not share a labeling rule, and the difference between the two rules is the subject of this paper.

##### Independent supervision (CRC 624)

Reaction activity is derived from the union of reactions present across 11 Human Metabolic Atlas reconstructions [12], projected onto Recon3D identifiers. Those 11 models are NCI-60 cancer cell-line reconstructions (786-O, HOP-62, HOP-92, HS-578T, HT29, MALME-3M, MDA-MB-231, NCI-H226, RPMI-8226, SR, UO-31), only one of which is colorectal; the union is our construction, not a released artifact, and identifier matching recovers between 2,140 and 2,690 reactions per model. Presence in a model is treated as active. The resulting vector marks 3,379 of 10,600 reactions active (31.9 %). Activity prevalence differs sharply between the two partitions: 40.8 % of expression-bearing reactions are labeled active, against 21.8 % of the remainder. Section 3.1 shows why that difference matters for every AUROC computed on the full set. Its virtue for our purpose is narrow and sufficient: it is computed from mass-balance reconstructions and not from the expression matrix supplied to the model. Its limitation is equally clear, and we return to it in Section 4.3.

##### Expression-derived supervision (BRCA, LUAD)

We state the rule exactly as implemented, because Section 3.2 turns on its details. For each reaction receiving nonzero expression, the cohort-mean value is compared against the 30th percentile of nonzero cohort means, and reactions active in fewer than 25 % of patients are forced inactive. For the remaining reactions, which receive no expression and admit no expression-derived score, the released implementation assigns labels by drawing from `numpy.random.RandomState(42)` with an activity probability of 0.70. The stated intent of that choice, recorded in the source comments, was to avoid a label pattern perfectly correlated with graph topology. Its consequence, which we did not recognize at the time, is that 4,675 of the 10,600 BRCA labels (44.1 %) carry no information about any patient or any reaction. The resulting vector is 68.7 % active. The first half of this rule follows the prevailing convention in the literature; the second half is our own error. We retain and audit both, because the audit is the contribution.

#### Gene identifier mapping

Projecting expression onto reactions requires bridging two identifier systems: the Recon3D GPR table uses Entrez identifiers with transcript suffixes (for example 8639.1) while the expression matrices are indexed by versioned Ensembl identifiers. We build the bridge through HUGO symbols, taking Ensembl → symbol from the STAR count annotation and symbol → Entrez from the Recon3D SBML gene-product records. On the CRC cohort this resolves 5,635 of the 5,938 GPR-bearing reactions (94.9 %).Section 3.4 reports why this step had to be rebuilt.

#### Splits and subsampling

All splits are patient-level. The archived CRC experiments draw a stratified subsample of 499 of the 624 patients per seed and apply 5-fold stratified cross-validation within it, giving folds of roughly 339 training, 60 validation and 100 test patients; ten seeds together cover all 624 patients but no single run saw more than 499. The verified retraining of Section 3.4 uses a 100-patient subsample and 3 folds. The cross-cancer cohorts use a single held-out test split. The reaction axis is never held out, so every reaction seen at test time was also seen during training. Section 3.2 shows that this is not a neutral design choice when the label vector is shared across patients.

#### Proteomics

The architecture reserves a proteomic feature channel, described in Module 3, but no CPTAC proteomic matrix could be aligned to the TCGA barcodes used here: no pre-computed quantitative matrix is available from PDC000111, and PDC000116 has no patient overlap with this cohort. The channel is retained and zero-filled for every patient, with an availability mask, rather than silently dropped. No result in this paper depends on proteomic input, and no claim of proteomic signal is made.

### 2.2 Module 2: Metabolic Network Graph Construction

#### 2.2.1 Heterogeneous Bipartite Graph Encoding

Figure 1 summarizes the pipeline and Figure 2 illustrates the graph encoding on a glycolysis subnetwork. Recon3D is encoded as a directed heterogeneous bipartite graph:

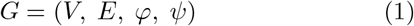

where the node set is the disjoint union of reaction and metabolite nodes:

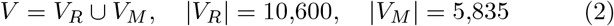

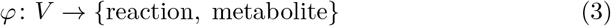

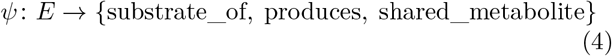

**Figure 1.**
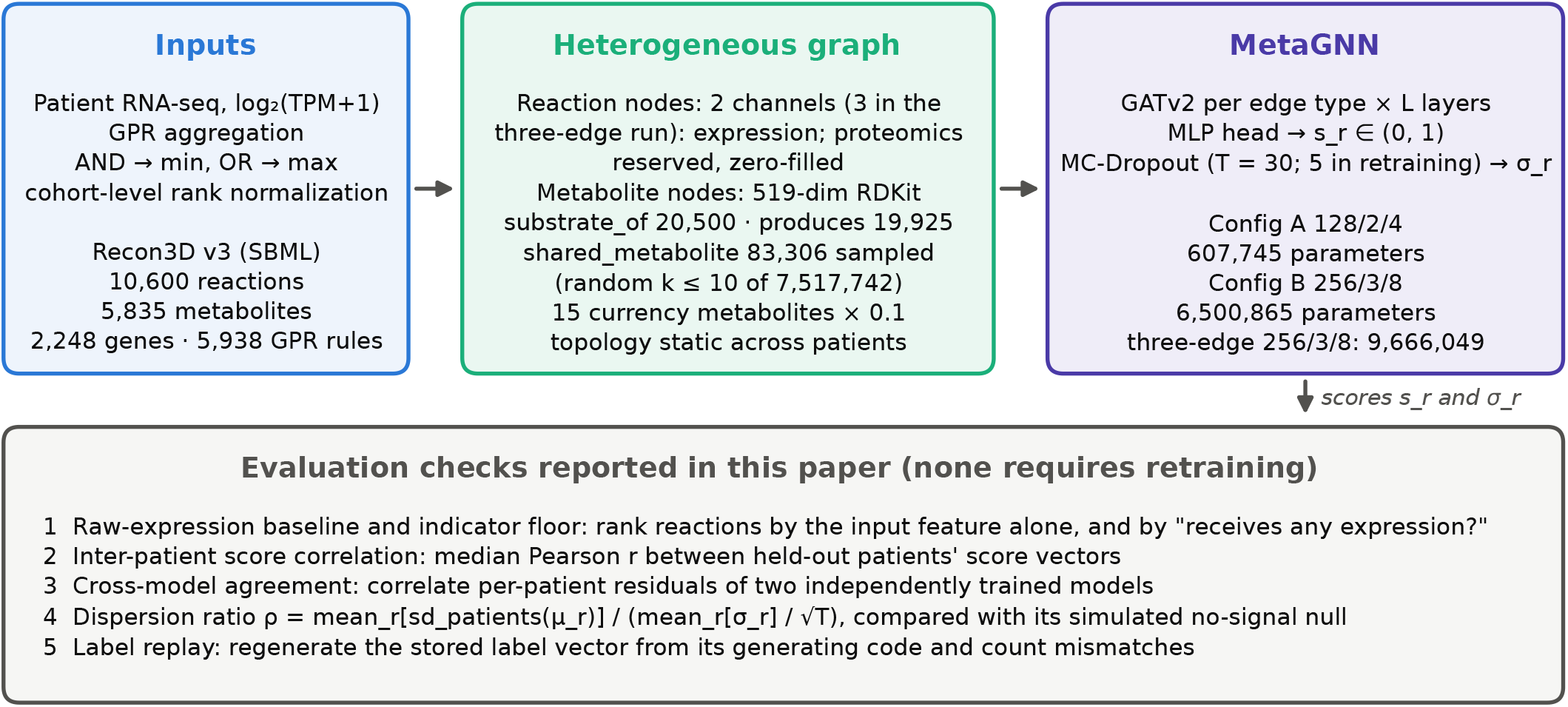
Pipeline and the checks applied to it. Patient expression is projected onto Recon3D reactions through GPR rules and propagated by GATv2 layers over a static heterogeneous graph; MC-Dropout supplies a per-reaction dispersion *σ*_*r*_ alongside the score *s*_*r*_. Node and edge counts are those of the parsed model used throughout; parameter counts are read from the released checkpoints. The lower panel lists the five evaluation checks of Section 2.7, all of which operate on artifacts a trained run already produces. No cohort size or split appears in this figure because none of the pipeline’s structure depends on the cohort.

**Figure 2.**
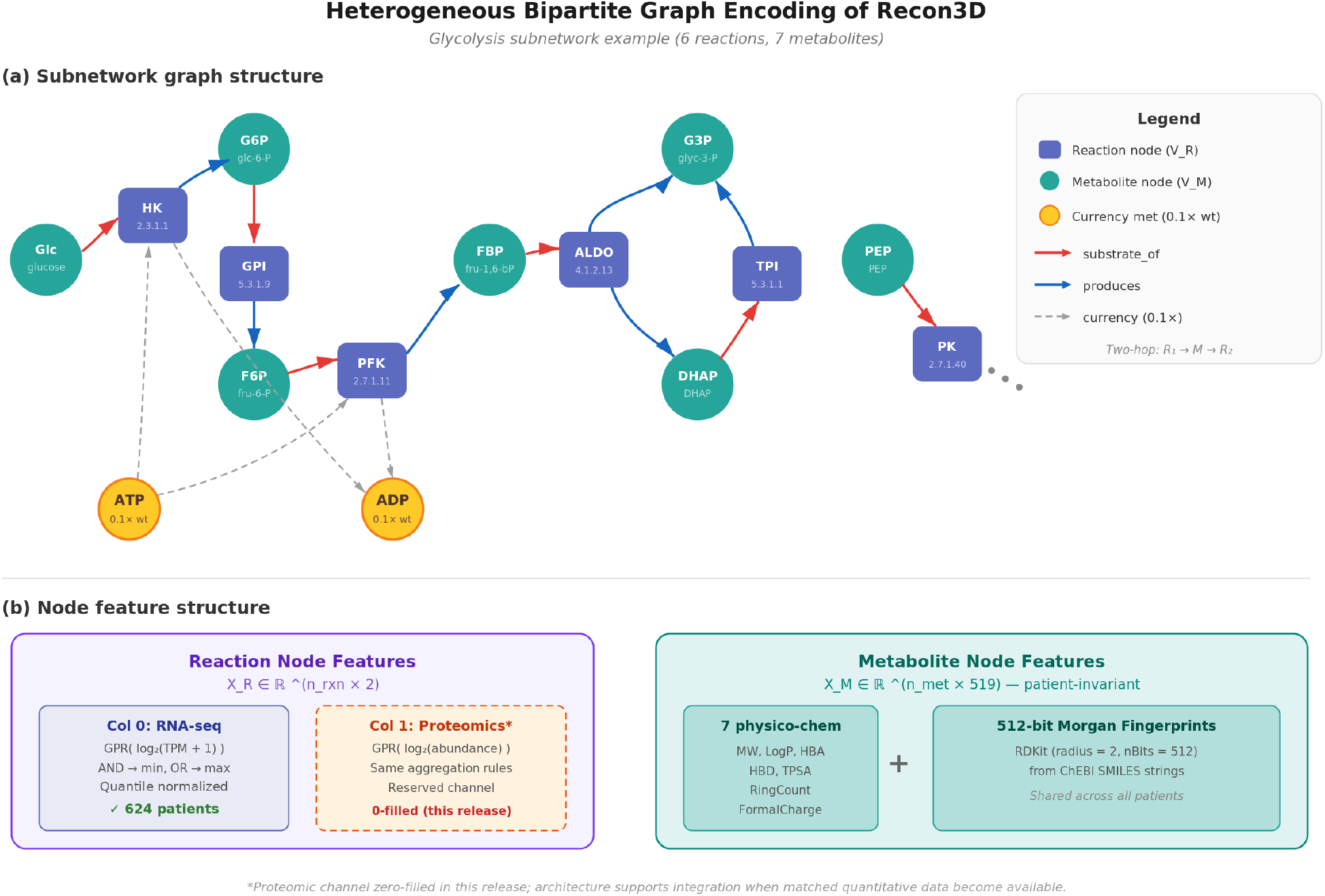
Heterogeneous bipartite encoding of Recon3D, illustrated on a glycolysis subnetwork. (**a**) Reaction nodes (squares) and metabolite nodes (circles) are joined by directed `substrate_of` and `produces` edges; the fifteen currency metabolites are down-weighted by a factor of 0.1; reaction-to-reaction communication is otherwise two-hop, through a shared metabolite. (**b**) Node feature structure: each reaction node carries a two-column vector (GPR-aggregated RNA-seq expression; a reserved proteomic column that is zero-filled in every release), and each metabolite node a patient-invariant 519-dimensional RDKit descriptor. Cohort sizes and splits are given in Section 2.1.

A directed edge (*m, r*) of type substrate_of exists whenever metabolite *m* is consumed by reaction *r* (stoichiometric coefficient *S*_*m,r*_ *<* 0), and an edge (*r, m*) of type produces exists whenever *r* produces *m* (*S*_*m,r*_ *>* 0). This encoding yields 20,500 substrate_of edges and 19,925 produces edges, for a total of 40,425 directed edges. Edge features encode the absolute stoichiometric coefficient |*S*_*m,r*_| and a binary reversibility indicator.

A third edge type, `shared_metabolite`, connects pairs of reactions that share at least one non-currency metabolite substrate or product, providing direct reaction-to-reaction information channels that supplement the indirect, two-hop connectivity available through the bipartite stoichiometric edges. On Recon3D, this relation generates 7,517,742 undirected edges (approximately 186× denser than the bipartite stoichiometric edges combined), reflecting the high interconnectivity of the metabolic network through shared intermediates. To make training tractable on consumer hardware (16–32 GB VRAM), we apply a random-*k* sparsification strategy: for each reaction node, *k* shared-metabolite neighbors are sampled uniformly at random and retained (we describe it as random rather than top-*k* because no ranking is applied), reducing the edge set to at most *k* × |*V*_*R*_| edges (fewer in practice because some reactions have fewer than *k* shared-metabolite neighbors). At the default *k* = 10, this yields 83,306 edges (a 99 % reduction from the full edge set), retaining a uniform random sample of each reaction’s metabolic neighborhood. We did not test how much neighborhood structure survives this reduction. The one experiment we ran with this edge type is reported, and withdrawn, in Section 3.3; we did not sweep *k*.

The graph structure is stored as a PyTorch Geometric [16] `HeteroData` object and loaded once at training time; the topology is static across patients, while node feature matrices vary per patient.

#### 2.2.2 Currency Metabolite Handling

We down-scaled edge weights by a factor of 0.1 during message aggregation for fifteen ubiquitous cofactors (ATP, ADP, AMP, NAD^+^, NADH, NADP^+^, NADPH, FAD, FADH_2_, CoA, H_2_O, H^+^, Pi, CO_2_, O_2_), to reduce without eliminating their influence. This is intended to limit spurious information pathways between otherwise unrelated reactions, a known artifact when currency metabolites are treated identically to pathway-specific substrates [17].

The fifteen species were selected by name from the standard cofactor list rather than by a degree cutoff. We report this precisely because an earlier description of this step claimed a degree threshold of 150, which does not in fact select this set. In our parsed Recon3D graph 31 metabolite nodes exceed degree 150, three of the fifteen named species never reach it (FAD and FADH_2_ at 148, CO_2_ at 107). Of the 31 nodes above 150, only 12 are among the fifteen, leaving 19 that are not cofactors in the intended sense (among them sodium, bicarbonate, carnitine, pyrophosphate and mitochondrial acetyl-CoA). The degree distribution shows no clean bimodal gap at that value. We did not run a sweep over the 0.1 scaling factor on the cohorts retained in this manuscript, so its contribution is uncharacterized and should be read as a design choice rather than a tuned parameter.

### 2.3 Module 3: Transcriptomic Feature Engineering (Proteomics-Ready)

#### 2.3.1 Transcriptomics

TCGA STAR-aligned gene counts (TPM values) were log_2_-transformed after adding a pseudocount of 1, then mapped to reaction nodes via GPR rules using a standard aggregation: AND relationships take the minimum across subunit genes, OR relationships take the maximum across isozymes (the convention stated by Gopalakrishnan et al. [9]). Values were normalized to (0, 1) by rank-based quantile transformation across the patient cohort.

#### 2.3.2 Proteomics

The pipeline reserves a second feature column for proteomic abundance, mapped to reaction nodes via GPR rules identically to transcriptomics. When both transcriptomic and proteomic data are available for a gene, a weighted average (*w* = 0.5 each) is used, as the two signals capture non-redundant information about gene product availability. In the current TCGA-CRC release, this column is zero-filled for all 624 patients because no pre-computed quantitative matrix is available from PDC000111 (the TCGA Retrospective Label Free study; Zhang et al. [18]), and PDC000116 (the CPTAC Prospective TMT study; Vasaikar et al. [19]) has zero patient overlap with the TCGA cohort (see Section 4.3). A binary mask tensor (`proteomics_available_mask.pt`) accompanies the release. For accuracy, its entries are all True, which is incorrect given that the proteomic column is zero for all 624 patients; the mask is not consumed by any code path used in this work, but it should not be relied upon.

#### 2.3.3 Metabolite Node Features

Metabolite nodes carry a 519-dimensional descriptor vector pre-computed using RDKit (v2023.09.5) from ChEBI SMILES strings: 7 physico-chemical properties (molecular weight, LogP, hydrogen bond acceptors, hydrogen bond donors, topological polar surface area, ring count, formal charge) and 512-bit Morgan fingerprints (radius 2, nBits = 512). These features are patient-invariant and provide structural and chemical similarity information for metabolite nodes during message passing.

#### 2.3.4 Patient Tensor Creation

The preprocessing step (the Module-3 driver together with `metagnn/data_loader.py`) produces one feature tensor per patient, containing the patient-specific reaction node features and shared metabolite features in the Hetero-Data format expected by the training script. For the TCGA-CRC cohort, this yields 624 patient tensors plus the shared graph structure file (one per unique patient after sample-level deduplication).

### 2.4 Module 4: Model Architecture and Training

#### 2.4.1 Architecture

The model backbone comprises stacked GATv2 layers (Brody et al. [10]) with type-specific weight matrices per edge type. GATv2 was chosen over the original GAT [20], itself a successor to spectral [21] and sampling-based [22] graph convolution, because GAT computes only *static* attention: the ranking of attention scores over a node’s neighbors is unconditioned on the query node. GATv2’s dynamic attention removes that restriction, so different reaction nodes can rank the same shared substrate differently, which is the behavior we require when substrate availability should modulate a reaction’s score. Each layer processes reaction and metabolite node types independently, with edges routed to the appropriate subgraph based on edge type. The output MLP head produces a per-reaction activity score *s*_*r*_ ∈ (0, 1) via sigmoid activation.

Two architecture configurations are provided, each of which can operate in either a *bipartite-only* mode (2 edge types: substrate_of, produces) or an *expanded* mode that additionally includes the `shared_metabolite` reaction– reaction edge type (3 edge types). Because GATv2 instantiates separate weight matrices per edge type, adding the third edge type increases the parameter count substantially.

Table 1 gives the parameter counts, read directly from the released checkpoints rather than derived analytically. An earlier version of this manuscript reported analytically estimated counts of 143,489 and 873,217; those figures were wrong in both magnitude and ordering, and we give the measured values here.

**Table 1.** Trainable parameter counts, summed over the tensors in the released checkpoints. The three-edge configuration adds a reaction → reaction GATv2 module per layer and a third input channel. Its 49 % parameter increase over the bipartite 256/3/8 configuration is one reason the comparison in Section 3.3 cannot isolate topology.

| Configuration | Parameters |
| --- | --- |
| Config A (128/2/4, bipartite) | 607,745 |
| Config B (256/3/8, bipartite) | 6,500,865 |
| Three-edge expanded (256/3/8) | 9,666,049 |

Both configurations are lightweight enough to train on consumer hardware (tested on Apple M4 Max with MPS acceleration and CPU fallback for unsupported operations).

#### 2.4.2 Expression-Derived Label Generation

Ground-truth reaction activity labels do not exist at scale for individual patients. For the two cross-cancer cohorts we used an expression-thresholding strategy, which is the prevailing convention in this literature. Section 2.1 states the rule as implemented, including the randomized assignment applied to reactions without expression. Two properties matter for interpreting the results. First, the threshold is computed from the same expression matrix supplied to the model, so the label for every expression-bearing reaction is a deterministic function of that input. Second, a single binary vector is shared by all patients in a cohort, so no patient-specific target exists.

An earlier revision of this pipeline used a per-gene cohort-median rule with majority voting across patients. It was abandoned during development because after rank normalization a large share of reaction scores tie exactly at the median, so strict inequality drove the consensus below one half for essentially every expression-bearing reaction, leaving a label set perfectly correlated with graph topology. The current rule was adopted to break that correlation, and Section 3.2 shows that it introduced a different and more consequential artifact.

For the 624-patient CRC cohort the labels come instead from the reconstruction union described in Section 2.1 and are independent of the expression input. An initial attempt to use the *generic*, non-tissue-specific HMA bounds was discarded: those bounds retain non-zero values for all 10,600 Recon3D reactions and yield a trivially all-active label set. Class weights inversely proportional to class frequency are applied during training in both regimes.

#### 2.4.3 Training Procedure

Training uses the AdamW optimizer with cosine annealing. Hyperparameters differ by cohort and we state them rather than averaging over them: the cross-cancer runs used learning rate 5 × 10^−4^, weight decay 10^−5^ and early-stopping patience 20; the CRC full-cohort runs used learning rate 10^−3^, patience 15, and weight decay 10^−5^ for the 128/2/4 configuration and 10^−4^ for 256/3/8; the verified retraining of Section 3.4 used learning rate 10^−3^ and patience 5. Dropout of 0.2 is applied within each GATv2 layer. The loss function combines weighted binary cross-entropy with a mass-balance regularization term:

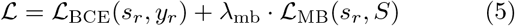

where ℒ_BCE_ is the standard binary cross-entropy between predicted scores *s*_*r*_ and pseudo-labels *y*_*r*_ (with class weights inversely proportional to class frequency), and the mass-balance term is:

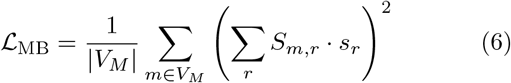

Because *s*_*r*_ ∈ (0, 1) is a predicted activity probability rather than a continuous flux value, *S*_*m,r*_ · *s*_*r*_ conflates stoichiometric coefficients with probabilities. The regularizer therefore acts as a *soft structural prior* encouraging metabolically coherent on/off patterns (balanced numbers of producing and consuming reactions per metabolite) rather than enforcing true mass-balance in the thermodynamic sense; enforcing exact *Sν* = 0 would require predicting continuous fluxes, which is outside MetaGNN’s current scope. With this caveat, the term penalizes predicted reaction sets where, for a given metabolite *m*, the weighted net flux ∑_*r*_*S*_*m,r*_ *s*_*r*_ deviates substantially from zero. The coefficient *λ*_mb_ = 0.2 was used throughout. We do not report a sweep over *λ*_mb_ on the cohorts retained in this manuscript, so the contribution of this term is uncharacterized here and should be treated as a design choice carried over from earlier development rather than a tuned or validated hyperparameter. Early stopping monitors validation F1 score in all cases.

### 2.5 Module 5: Uncertainty-Aware Inference

Monte Carlo Dropout [11] is applied at test time to estimate per-reaction epistemic uncertainty. Dropout layers remain active during *T* stochastic forward passes (*T* = 30 for the archived runs, *T* = 5 for the reduced-scale retraining), yielding a distribution of predicted scores 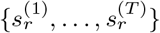 for each reaction *r*. The predictive mean and standard deviation are:
xsxs

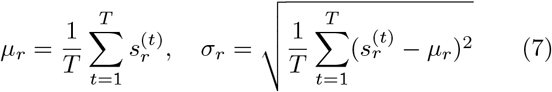

We use *σ*_*r*_ for two purposes only: selecting boundary reactions for the module of Section 2.6, and forming the denominator of the dispersion ratio defined in Section 2.7. We attach no probabilistic interpretation to it.

### 2.6 Module 6: Optional LLM Boundary Validation

Monte Carlo Dropout identifies reactions on which the model is least decided. We built an optional post-hoc module that routes those reactions to a debate-and-reconcile loop over locally hosted open-weight language models, and report it here because it was part of the pipeline evaluated in Section 3.6, not because its results support a claim.

#### Boundary selection

For each patient, reactions with MC-Dropout predictive mean in [0.3, 0.7] are ranked by predictive standard deviation and the top 500 are retained per fold. On the archived CRC runs this pool is 0.08 ± 0.06 % of reactions for the 128/2/4 configuration and 11.31 ± 5.03 % for 256/3/8 (means over ten seeds), corresponding to medians of roughly 7 and 1,125 reactions respectively. The gap is a property of output dispersion, not of accuracy, and Section 3.3 shows the underlying scores were not patient-specific.

#### Loop structure

Figure 3 shows the loop. The module is implemented as a LangGraph [23] StateGraph with three sequential computation nodes: two advocate calls that argue for and against activity from the assembled evidence, and one resolver call that adjudicates and returns a calibrated score with a rationale. Evidence for each reaction comprises its Recon3D record, KEGG pathway context where retrievable, passages retrieved from a FAISS index over a curated corpus of reaction descriptions and enzyme summaries using an `all-MiniLM-L6-v2` sentence-transformer, and subsystem-level context for reactions without a GPR association. The final score combines the GNN and resolver outputs as *s*_final_ = *w*_GNN_*µ*_*r*_ +*w*_LLM_*s*_LLM_ with *w*_GNN_ = 0.7 and *w*_LLM_ = 0.3, applied only when the resolver returns a confidence above 0.8; otherwise the GNN score is retained unchanged. Sampling temperature is 0.1 throughout, and the state object carries 46 typed fields recording every input and intermediate so that runs are auditable after the fact.

**Figure 3.**
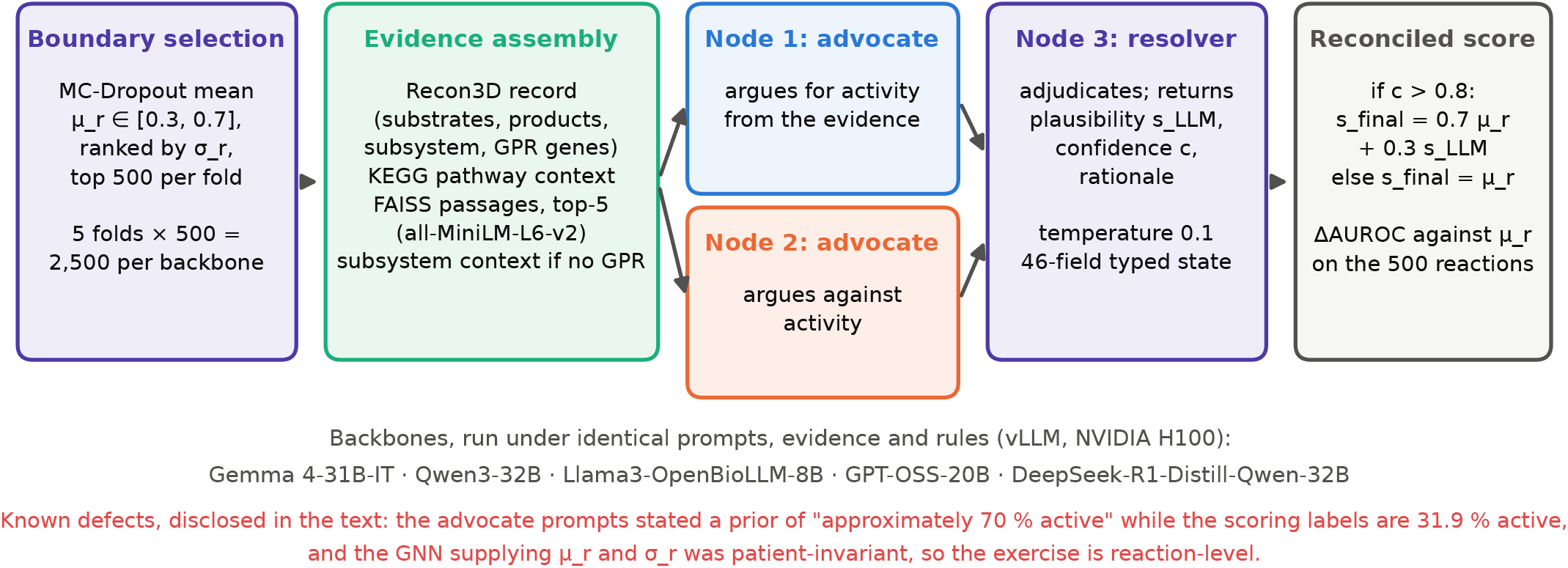
Optional LLM boundary-validation module. Reactions on which the GNN is least decided are routed, with assembled evidence, through a three-node debate-and-reconcile graph; the reconciled score replaces the GNN score only when the resolver reports confidence above 0.8. Five open-weight backbones were run under identical conditions (Section 3.6). The two defects noted in red are discussed in Section 3.6 and are the reason the module’s results are reported as descriptive only.

#### Backbone comparison

Five open-weight backbones were run under identical prompts, evidence and decision rules, five folds of 500 boundary reactions each, served through vLLM [24] on NVIDIA H100 80 GB hardware, with tensor parallelism across two devices for the two largest models and a single device otherwise. Section 3.6 reports the outcome. We designed this as a cumulative ablation over six module versions; only the final version was carried through to the backbone comparison, and we report no results for the intermediate versions because none was evaluated against held-out labels.

### 2.7 Evaluation Strategy

Five checks anchor the results below. Each is cheap and each is computable from artifacts a normal run already produces. We introduce them together because the failures reported in Section 3.2 and Section 3.3 are of different kinds and no single check catches both.

#### 1. The raw-expression baseline, and an information-free floor beside it

We compute the AUROC obtained by ranking reactions using the GPR-mapped expression value alone, with no model. Because the expression-derived labels are defined from the *cohort-mean* feature, we report that ranking as well as the per-patient ranking, and say which is which at each use. We also report the AUROC of a binary indicator of whether a reaction receives any expression at all. That indicator contains no transcriptomic information, so the gap between it and the expression baseline is the part of the baseline that expression actually supplies. Reporting the indicator matters because activity prevalence differs systematically between the two partitions, and a full-set AUROC therefore carries a component that any method inherits for free.

#### 2. The inter-patient score correlation

We report the median Pearson correlation between per-reaction score vectors across pairs of held-out patients. A high value shows that the model’s output is nearly the same for every patient. We record in advance what this check can and cannot do: it detects that a model has collapsed onto a shared vector, which is the failure of Section 3.2, but it does not distinguish that case from a model given no patient input at all, because both produce near-identical outputs. In our results it returns values above 0.99 for every model in this study, including the one that is reading its input.

#### 3. Cross-model agreement on patient deviations

For two models trained independently on the same cohort, we center each score matrix per reaction and correlate the residuals across the patients they share. Models that read patient input must agree, at least weakly, on which patients deviate in which direction; models that do not read it have nothing to agree about. This check needs two runs and no ground truth.

#### 4. The dispersion ratio

Monte Carlo Dropout gives each prediction a sampling standard deviation *σ*_*r*_ over *T* stochastic passes, so the sampling standard error of a predictive mean is 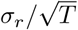. If patients differ only by dropout randomness, the between-patient standard deviation of the predictive means will match that quantity. We therefore report

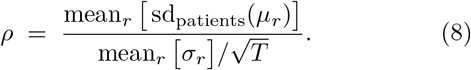

This is a ratio of standard deviations, not of variances. Its no-signal value is not exactly 1: both pipelines compute *σ*_*r*_ and the between-patient spread with a 1*/n* normalization, which biases the ratio upward, and simulating the null with the observed *σ* field gives 1.019 at *P* = 100 patients and 1.023 at *P* = 165. We therefore compare each measured *ρ* against that simulated null rather than against 1. Two caveats are structural. *ρ* is normalized by each model’s own dropout magnitude, so it measures patient signal relative to that model’s noise and is not directly comparable across models with very different *σ*. A value at the null is consistent with input invariance but does not prove it, since a real signal much smaller than the dropout noise would also land there. It is a screening check, and in this paper it is corroborative rather than load-bearing: the conclusion of Section 3.3 rests on the feature tensors themselves and on check 3.

#### 5. Label replay

Where a label vector was produced by code, we regenerate it from that code and count mismatches against the stored vector. This is the only check here that examines the target rather than the model, and it is the one that found the randomized partition of Section 3.2.

##### Metrics

We report AUROC as the primary threshold-free ranking metric, and positive-class F1 at a fixed decision threshold: 0.5 for the cross-cancer cohorts and 0.15 for the CRC cohort, in both cases sklearn’s default binary averaging as implemented in the released evaluation scripts. An earlier version of this manuscript described the CRC figures as Macro-F1; they are not, and macro averaging on the same predictions would give different values. AUROC presents an optimistic picture on imbalanced targets relative to precision–recall analysis [25]; we retain it because it is the metric this literature reports and because the effects we document are far larger than the metric choice. Because a large minority of reactions receive no expression signal (4,675 of 10,600 on BRCA, 4,965 on the rebuilt CRC features, against a nominal 4,662 without a GPR rule), we report the expression-bearing and non-expression partitions alongside full-set figures wherever both are available, and we flag where they are not. We report no calibration metric; MC-Dropout outputs should be read as uncertainty-aware rather than calibrated, and we use their dispersion only to select boundary reactions and to compute *ρ*, never as a probability statement.

##### Statistics

Experiments with stochastic components report mean ± standard deviation across the stated replication unit (seeds, folds or patients), with the unit named at each use. Confidence intervals are bootstrap percentile intervals over the stated unit with 2,000 resamples. We use 5-fold stratified cross-validation to bound compute cost; Kohavi [26] recommends 10 folds, and the reduction is ours. Standard deviations use the 1*/n* normalization throughout, matching the released evaluation code. Group comparisons use the Mann–Whitney *U* test. Where a ± figure appears without a named unit in a table, the unit is given in that table’s caption. Several comparisons are reported, so we interpret individual *p*-values descriptively and apply no family-wise correction.

### 3 Results

#### 3.1 The raw-expression baseline on independent supervision

Because the 624-patient CRC labels derive from HMA reconstructions rather than from the expression matrix, this cohort admits a question the cross-cancer settings cannot answer: does graph-based scoring extract signal beyond what the input feature already carries?

Averaged over all 624 patients, ranking reactions by GPR-mapped expression alone attains AUROC 0.6342 ± 0.0058 over the full 10,600-reaction set (bootstrap 95 % confidence interval over patients [0.6338, 0.6347]) and 0.5755 ± 0.0177 over the 5,635 expression-bearing reactions. Per-patient values span 0.6185 to 0.6540, so the baseline is stable across the cohort rather than an average over a wide spread.

An information-free floor sits close underneath it. Ranking reactions by a binary indicator of whether they receive any expression at all, with no transcriptomic values used, gives AUROC 0.6085 on the same labels, because activity prevalence differs between the two partitions (40.8 % against 21.8 %). Expression values therefore contribute roughly 0.026 of the 0.6342; the rest is prevalence structure that any method inherits without doing anything. On the expression-bearing partition alone, where that structure is absent, the raw-expression baseline is 0.5755 ± 0.0177.

We report both numbers because the full-set figure is the one a model is scored against, and the indicator is what that figure is worth. One further caveat on comparability: the baseline is averaged over all 624 patients, whereas the model result of Section 3.4 is a held-out mean over a 100-patient subsample, and the two ± figures are dispersions over different units.

#### 3.2 Under expression-derived supervision: what the near-ceiling figure is made of

Applying the pipeline to TCGA-BRCA gives AUROC 0.9864 ± 0.0008 across 165 held-out patients; TCGA-LUAD gives 0.9839 ± 0.0005 across 79. The partition analysis below is performed on BRCA, for which the per-patient score archive was retained; LUAD was generated by the identical code path and we expect it to behave the same way, but we have not verified that and do not claim it. Partitioning the BRCA reactions by whether they receive expression identifies what each part contributes (Table 2, Figure 4).

**Table 2.** TCGA-BRCA test AUROC by reaction partition, computed per patient over the 165 held-out patients from the released score archive. Neither component measures metabolic biology. The partition AUROCs do not average to the full-set value, because AUROC over a pooled set also counts cross-partition pairs; the partitions are reported to identify what each contributes, not as an arithmetic decomposition.

| Partition | Reactions | AUROC |
| --- | --- | --- |
| All reactions | 10,600 | $0.9864 \pm 0.0008$ |
| Expression-bearing | 5,925 | $1.0000 \pm 0.0000$ |
| No expression | 4,675 | $0.9291 \pm 0.0030$ |

**Figure 4.**
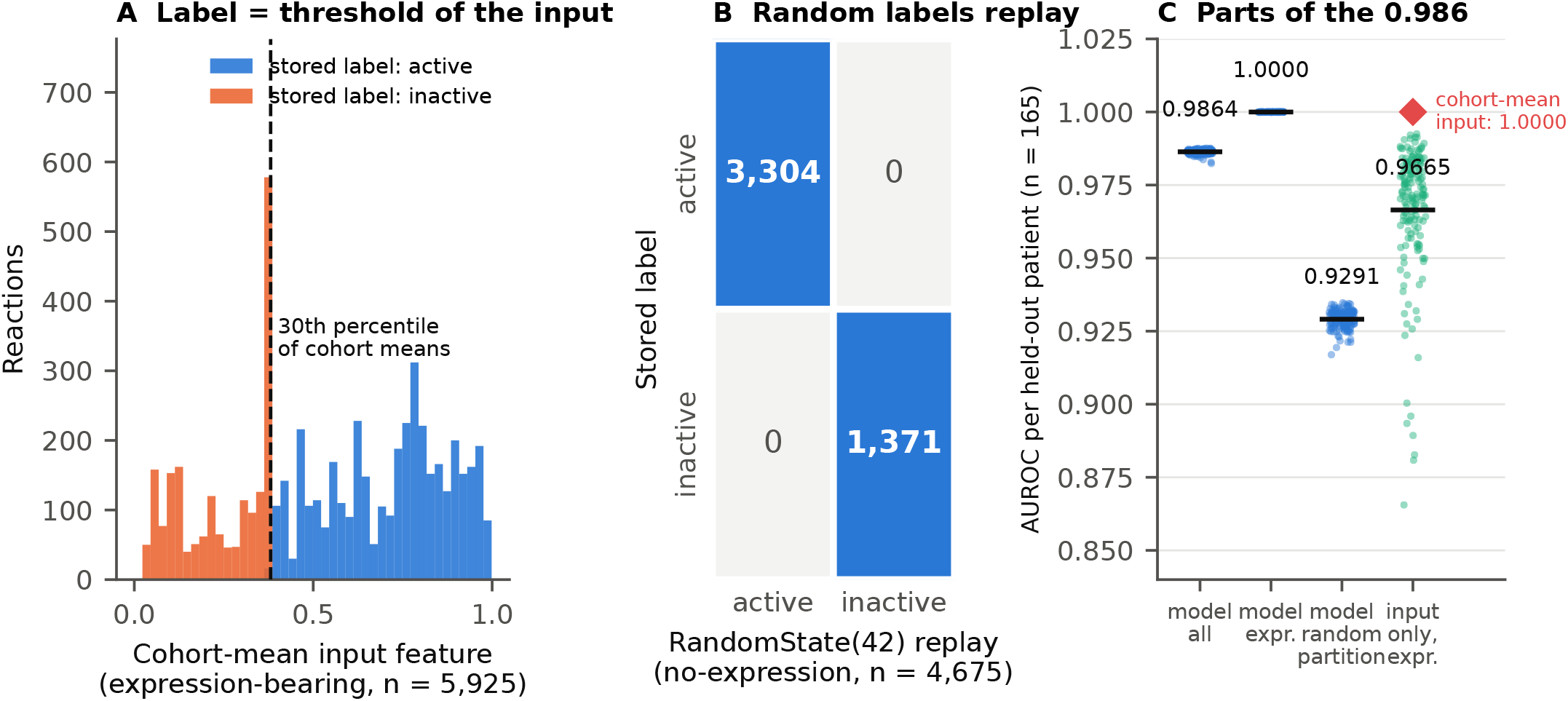
What the TCGA-BRCA benchmark score is made of. (**A**) For the 5,925 expression-bearing reactions, the stored activity label is exactly the cohort-mean input feature thresholded at its 30th percentile; the 25 % prevalence rule in the released code affected no reaction on this cohort. (**B**) For the 4,675 reactions receiving no expression, the stored labels (3,304 active, 1,371 inactive) are reproduced with zero mismatches by `numpy.random.RandomState(42).random(4675)` < 0.70. (**C**) Per-patient AUROC over the 165 held-out patients: the model reaches 1.0000 on the partition whose label is a function of its input, 0.9291 on the partition whose labels are random, and 0.9864 overall; ranking each patient’s own input feature, with no model, reaches 0.9665 on the expression-bearing partition (green points), and ranking the cohort-mean feature reaches 1.0000 exactly (red diamond). Bars mark means.

#### The expression-bearing partition is the model reading back its own input

On these 5,925 reactions the label is, by construction, a threshold of the cohort-mean input feature. Ranking by that same cohort-mean feature reproduces the label set at AUROC 1.000000; ranking by an individual patient’s feature gives 0.9665 ± 0.0228 over the 165 test patients. The gap between those two numbers is the entire patient-level content of this partition, and the model’s 1.0000 ± 0.0000 sits at the cohort-mean value rather than the per-patient one. A model given that feature and scored on that label is being measured on how completely it reconstructs a cohort-level transformation of its own input.

#### The remaining partition is memorization of noise

These 4,675 reactions receive no expression, so their stored labels were assigned by a seeded pseudo-random generator. We confirmed this directly: drawing 4,675 values from `numpy.random.RandomState(42)` and thresholding at 0.70 reproduces the stored labels for exactly these reactions, with zero mismatches. The labels are independent of every patient, every gene and every network property.

A model cannot generalize to such labels. It can memorize them, and the AUROC of 0.9291 ± 0.0030 is what that memorization records. The mechanism is structural rather than accidental: the label vector is shared by all patients and the splits are patient-level, so every reaction’s label appears in the training set of every fold. Held-out patients present the model with the same 10,600 reactions it was fit on, and a per-reaction bias learned during training transfers to them intact. Standard patient-level cross-validation offers no protection, because nothing on the reaction axis is ever held out. This is a variant of a failure mode documented elsewhere in computational biology, where structure shared between training and evaluation partitions inflates cross-validation estimates [27], and it is consistent with reports that zero-shot evaluation of single-cell foundation models can leave them outperformed by simpler methods [28].

#### The outputs are near patient-invariant

Across the 165 BRCA test patients the median inter-patient score correlation is 0.995, with a mean between-patient standard deviation of 0.019 per reaction. On the no-expression partition alone the median is 0.986, which is what a memorized constant vector looks like. We note against our own interest that this diagnostic does not discriminate here: the three colorectal models of Section 3.3, which received no patient input at all, return medians of 0.9986, 0.9959 and 0.9953, so a high inter-patient correlation is consistent with both a model collapsed onto a shared label and a model with no input. It tells you something is wrong, not which thing.

#### Orthogonal biological checks are weak

Mapping the 10,600 reactions to 106 Recon3D subsystems and ranking by mean predicted activity, 3 of 8 literature-derived breast cancer expectations are concordant. Glutamate metabolism ranks 14th of 106 and folate metabolism 22nd, both consistent with elevation; squalene and cholesterol synthesis ranks 103rd, consistent with reduction. The other five are not: glycolysis ranks 91st despite the expected Warburg elevation, purine synthesis 90th, pyrimidine synthesis 98th, fatty acid synthesis 42nd, and oxidative phosphorylation 50th where reduction was expected. On LUAD, concordance is 0 of 5. The expectation lists are given in the deposited analysis script.

#### Against analytical baselines

On the same expression-thresholded target and a 50-patient evaluation subset, MetaGNN reaches 1.000 on expression-bearing reactions where GIMME-style scoring reaches 0.962 ± 0.023 on BRCA and 0.970 ± 0.016 on LUAD, iMAT-style scoring reaches 0.820 and 0.813, and a simple expression threshold reaches 0.660 and 0.679. All four figures are computed on the GPR partition, so they are directly comparable with each other; on the full 10,600-reaction set the same threshold rule gives 0.575 and 0.598. We report these for completeness and not as a ranking. On this partition the label is a deterministic function of the cohort-mean input, so the comparison measures only which rule aggregates most faithfully to that transform. A method could top this table while carrying no biological information whatever, which is precisely what the rest of this section argues ours does.

We therefore report the BRCA and LUAD figures as within-cohort self-consistency scores on a shared, partly randomized target, and not as reconstruction accuracy or as evidence of patient-specific generalization.

### 3.3 The archived CRC models did not use their input

The cross-cancer failure above is a property of the labels. The CRC cohort has input-independent labels and should have been immune. It was not, for an unrelated reason.

While preparing the data deposit we found that the released per-reaction feature tensors for the 624-patient cohort are uniformly zero: across 624 files and 13.2 million entries there is no nonzero value. That directory is the `data_root` recorded in the run configuration of the two archived bipartite CRC experiments. The three-edge experiment overrides it with a separate feature directory that was never transferred off the training machine and is not in the deposit, so for that run the input cannot be inspected at all.

Three lines of evidence establish that none of the three received patient-specific input.

#### The feature files

Figure 5 summarizes the two quantitative checks. For the two bipartite runs this is direct: the model was given a matrix of zeros for every patient, so its forward pass cannot depend on patient identity.

**Figure 5.**
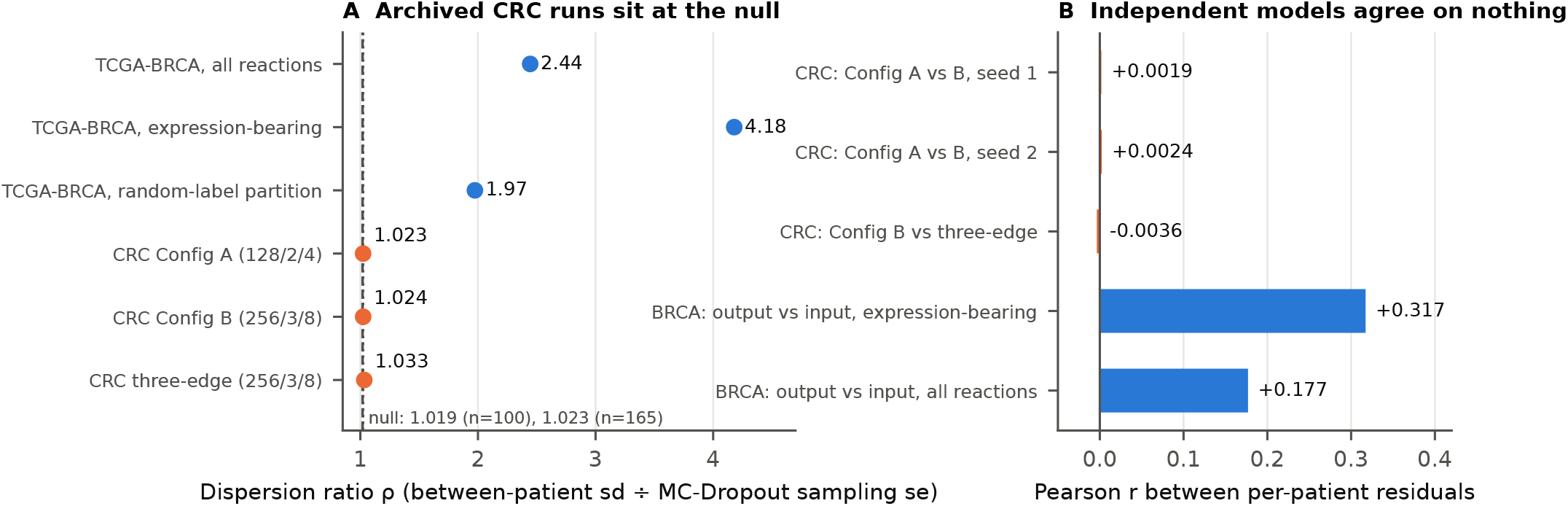
The archived CRC models did not use their input. (**A**) Dispersion ratio *ρ* for the three archived TCGA-CRC configurations (means over 20 seed–fold combinations; standard deviations, 0.001–0.003, are smaller than the markers) and for the TCGA-BRCA run with verified per-patient features, shown for all reactions and for its two partitions. The dashed and dotted lines mark the simulated no-signal null at 100 patients (1.019, the CRC runs) and 165 patients (1.023, BRCA). (**B**) Correlation between per-patient output residuals of independently trained CRC models, and, for comparison, between the BRCA model’s per-patient output residuals and its per-patient input residuals.

#### Independently trained models agree on nothing

Centering each score matrix per reaction and correlating the residuals across shared patients, models trained with different capacities and different seeds show no agreement on which patient deviates in which direction: *r* = +0.0019 between the two bipartite configurations at seed 1, +0.0024 at seed 2, and −0.0036 between Config B and the three-edge run. For comparison, on TCGA-BRCA the model’s per-patient output residual correlates with its per-patient input residual at *r* = +0.317 on expression-bearing reactions and +0.177 overall. This check needs no ground truth and no assumption about dropout, and it covers the three-edge run whose features we cannot inspect (Figure 5B).

#### The dispersion ratio

Table 3 reports *ρ* as defined in Section 2.7, together with the simulated no-signal null for the relevant patient count.

**Table 3.** Dispersion ratio *ρ* on archived per-patient predictions, with *T* = 30 passes. Values for the three CRC configurations are means ± standard deviations over 20 seed–fold combinations (archived seeds 1, 2, 4 and 8, folds 0–4; the archive holds ten seeds and we evaluated the first four); the BRCA row is a single held-out split. “Null” is the value *ρ* takes when patients differ only by dropout randomness, simulated with each setting’s own *σ* field and patient count. All four rows use the same *σ* estimator and the same *T*. The BRCA row is a sensitivity control, not a matched one: it differs from the CRC rows in cohort, label rule, architecture and topology, and shows only that *ρ* is capable of rising above the null when a model reads patient input.

| Run | Input features | Patients | Null | $\rho$ |
| --- | --- | --- | --- | --- |
| TCGA-BRCA full cohort | verified nonzero | 165 | 1.023 | **2.44** |
| CRC Config A (128/2/4, bipartite) | released tensors all zero | 100 | 1.019 | $1.0234 \pm 0.0017$ |
| CRC Config B (256/3/8, bipartite) | released tensors all zero | 100 | 1.019 | $1.0239 \pm 0.0028$ |
| CRC three-edge expanded (256/3/8) | override set, not archived | 100 | 1.019 | $1.0333 \pm 0.0021$ |

All three CRC configurations sit within 0.02 of the null in every seed and fold (maximum 1.0376), while the BRCA control sits at more than twice it (Figure 5A). We do not read the small ordering among the three CRC values as meaningful; they are separated by less than the bias the estimator itself introduces. Partitioning the BRCA control is instructive about what *ρ* measures: it is 4.18 on expression-bearing reactions and 1.97 on the memorized random partition, so a value well above the null indicates that patient input reaches the output, not that the output is useful.

#### What this invalidates

Three results we had previously reported rest on those runs and are withdrawn.

- The comparison between architecture configurations (AUROC 0.5437 ± 0.0051 for 128/2/4 and 0.5584 ± 0.0074 for 256/3/8 over ten seeds) is not a comparison of scoring accuracy. Neither model received patient input, so both figures describe how well a fixed output vector matches a fixed label vector under two capacities. These two runs also differ in weight decay (10^−5^ against 10^−4^) and epoch budget (80 against 200) as well as capacity, so they would not have isolated capacity even had the inputs been valid.
- The +0.105 AUROC gain from adding reaction– reaction shared-metabolite edges, which we had described as the largest single architectural lever, was the difference between 0.6637 ± 0.0025 for that run and 0.5584 ± 0.0074 for the bipartite 256/3/8 configuration. It came from a run that simultaneously changed the edge topology, substituted a three-channel feature set for the two-channel one, and raised the parameter count from 6,500,865 to 9,666,049. The gain was reported in an earlier, unposted draft of this manuscript and in the first version of the data deposit. That feature set was never archived, so the run cannot be reproduced or inspected, and it shows no patient signal on either of the two checks that do not require the features. We withdraw the relational-edge claim. It may still be true; this experiment does not bear on it.
- The microsatellite and clustering analyses were computed from the same predictions. Positive-class F1 was 0.4615 ± 0.0102 on MSI-H and 0.4610 ± 0.0093 on MSS patients (Δ = +0.0005; Mann–Whitney *U* = 22,949, *p* = 0.71, *n* = 92 and 487), and *k*-means with *k* = 2 on the seed-averaged scores gave a silhouette coefficient of 0.171. Neither is informative about biology as originally framed. The absence of MSI stratification is what an input-invariant model predicts; the silhouette value is not, and we report it without an account of it.

#### Why standard practice did not catch this

The archived runs used stratified patient-level cross-validation, ten random seeds, an inner validation split, early stopping on validation F1, and reported mean ± standard deviation. Every one of those controls behaved normally, because none examines whether a model’s output depends on its input. Multi-seed replication in particular produced tight standard deviations across seeds, which reads as stability and is equally consistent with a model that has nothing to vary. We want to be precise about the sequence, because it matters for what we can claim. The defect was found by inspecting the deposited files, not by a diagnostic, and the checks reported above were run afterward to establish its extent and to cover the run whose files are missing. What we take from that is not that these checks would have caught it first, but that they are cheap enough to run on every result, and that nothing else we ran would have caught it at all.

### 3.4 Retraining on verified features

We rebuilt the CRC feature tensors through the Ensembl → symbol → Entrez chain described in Section 2.1, resolving 5,635 of 5,938 GPR-bearing reactions (94.9 %) and confirming nonzero, patient-varying values for all 624 patients.

Retraining at reduced scale on CPU (100 patients stratified by project, 3-fold cross-validation, hidden dimension 64, two layers, four heads, three-edge topology, learning rate 10^−3^, patience 5, MC-Dropout *T* = 5) gives a held-out AUROC of 0.5800 ± 0.0017 across the three folds (0.5788, 0.5824, 0.5787). Two variants run on the first fold give 0.5815 with an extended epoch budget, early-stopped at epoch 25, and 0.5808 at doubled width. The three first-fold values span 0.0027, and doubling width moved that fold by 0.0020. This is consistent with convergence at this scale but does not establish it, since the variants share a fold and no learning curve was recorded.

This is the only CRC figure in the paper computed on features we have verified to be nonzero and patient-specific (Figure 6). It sits 0.054 below the raw-expression baseline of 0.6342 and 0.029 below the information-free indicator of 0.6085, though the comparison is across different patient sets and replication units, as noted in Section 3.1. We did not compute the expression-bearing and non-expression partitions of this figure, which is the omission a reader should press hardest on given the argument of Section 3.1. Whether a full-capacity model on the complete cohort would close the gap is untested. What we can say is that the archived numbers do not answer it.

**Figure 6.**
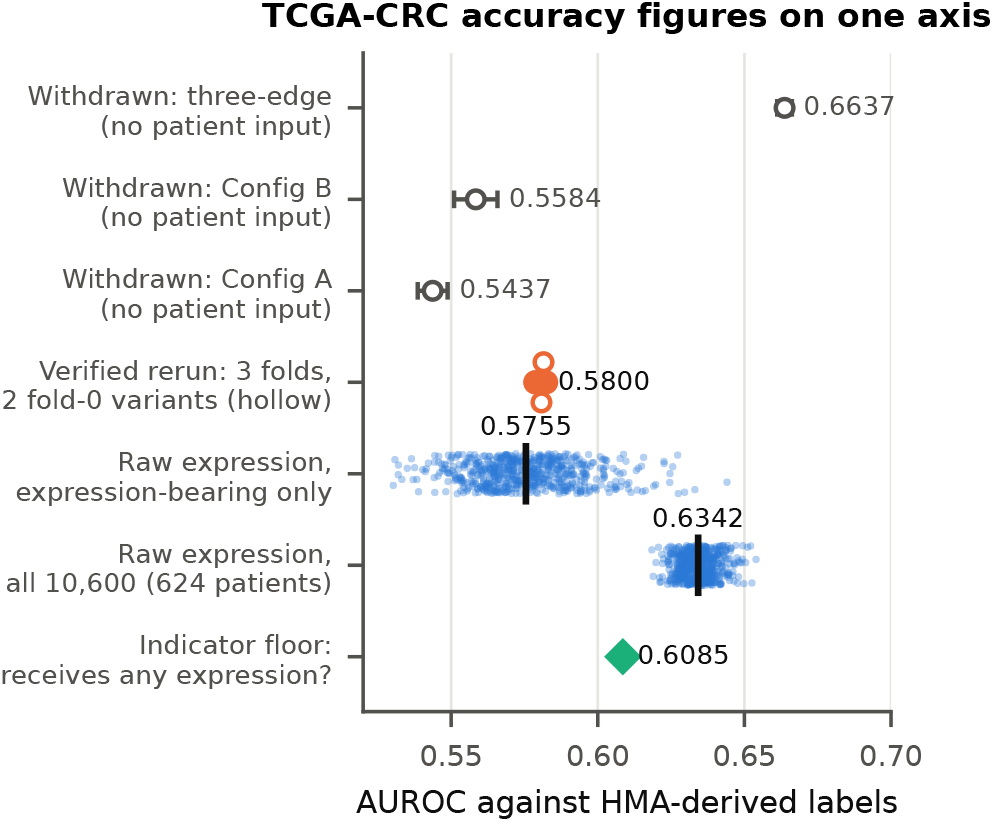
Every TCGA-CRC accuracy figure in this paper, on one axis. AUROC against the HMA-derived labels. Points for the two raw-expression rows are the 624 individual patients, with the mean marked; the verified rerun shows its three folds (filled) and the two single-fold variants (hollow), and the printed value is the 3-fold mean; the withdrawn archived runs are drawn hollow with their ten-seed standard deviations. The indicator floor uses no expression values at all.

### 3.5 Cross-consortium transfer and four discriminators

Zero-shot application of the BRCA-trained model to METABRIC gives AUROC 0.4926 ± 0.0113 over all reactions and 0.4913 ± 0.0222 on expression-bearing reactions (*n* = 200; gene coverage 1,795 of the 1,889 Recon3D genes appearing in a resolved GPR rule, or %). The confidence interval on the first figure excludes 0.5 from below, so this is at or marginally below chance rather than exactly at it. Four experiments, specified before they were run but not pre-registered, weigh candidate explanations (Figure 7).

**Figure 7.**
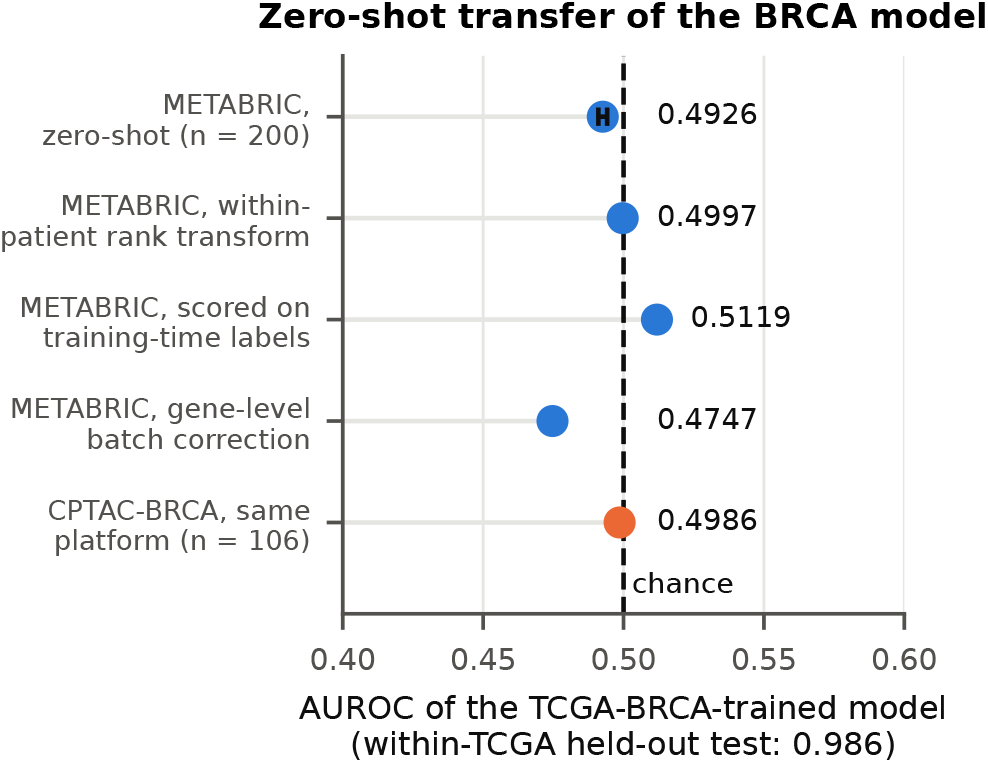
Zero-shot transfer of the BRCA-trained model. AUROC on METABRIC under the baseline evaluation and three of the four discriminators, and on CPTAC-BRCA, which shares the training platform. The error bar on the zero-shot value is the 95 % confidence interval of the mean over the 200 METABRIC patients, which excludes 0.5; the other discriminator values are single evaluations.

- **Rank transform**. Replacing the cohort-level rank normalization of Section 2.3.1 with a within-patient rank-to-quantile transform, to remove platform-specific distributional shape, moves AUROC from 0.4926 to 0.4997 (Δ+0.007 on all reactions, −0.001 on expression-bearing). Distributional shape is not the mechanism.
- **Training-time labels**. Re-evaluating the same predictions against the labels used at training time rather than METABRIC-derived labels gives 0.5119 (Δ+0.019 on all reactions, +0.038 on expression-bearing). Label construction accounts for a small fraction at most.
- **Gene-level batch correction**. Applying an empirical-Bayes style correction [29] across TCGA-BRCA and METABRIC on their 19,451 shared genes moves AUROC from 0.4924 to 0.4747, and from 0.4909 to 0.4640 on expression-bearing reactions. Correction makes transfer worse. This discriminator is the weakest of the four: our implementation applied a simplified mean-and-variance adjustment rather than the full empirical-Bayes estimator, so it constrains rather than excludes a per-gene batch mechanism. The small numerical differences from the baseline in items 1 and 2 reflect the gene intersection used for this experiment.
- **Same-platform control**. The most informative test. Applying the same model to CPTAC-BRCA, an RNA-seq cohort on the same assay platform as the training data, gives AUROC 0.4986 (*n* = 106), with a boundary pool of 0.017 %. Transfer fails even with the platform held constant.

Taken together these weigh against platform distribution and label construction as the dominant mechanism, and constrain without excluding a per-gene batch mechanism. We had originally read this as evidence for a consortium-specific representation.

We no longer think the evidence supports naming a mechanism. Section 3.2 offers a competing account, that a model which has collapsed onto a memorized label vector will score near chance against any differently constructed label set on any cohort, but discriminator 2 argues against the strong form of it. If the output were simply the memorized training-time vector, scoring it against that same vector would return a figure near the within-cohort 0.986, not 0.5119. The output does change with the input cohort; it simply does not change usefully. We report the four discriminators as constraints on the explanation and leave the explanation open.

### 3.6 LLM-based boundary validation

An optional post-hoc module routes the most uncertain reactions, those with MC-Dropout scores in [0.3, 0.7], to a debate-and-reconcile loop over locally hosted open-weight models. We evaluated five backbones under identical conditions, five folds of 500 boundary reactions each, drawn from the 624-patient CRC cohort. Because Section 3.3 shows that the model supplying those scores had no patient input, the exercise should be read as reaction-level rather than patient-level, and its starting point is chance by construction.

Two of five backbones produced a positive mean ΔAUROC (Table 4); one produced a clearly negative one. Both positives are instruction-tuned generalists, but so is GPT-OSS-20B, which did not improve, so instruction tuning does not separate the groups and we offer no mechanism. With five backbones, one cohort and a starting point at chance, this is a descriptive observation. Absolute post-reconciliation AUROC remains at or below 0.56 in every case, so at most this module improves a chance-level signal on the hardest subset of reactions, and it does so on scores that Section 3.3 shows were not patient-specific. One further defect should be recorded: the advocate prompts stated a prior of approximately 70 % active reactions for this cohort, a figure inherited from the expression-thresholded convention, whereas the HMA labels against which the module was scored are 31.9 % active. This is also consistent with reports that language models handle biological pathway reasoning poorly, particularly in perturbed systems [36]. We report the module for completeness and draw no conclusion from it.

**Table 4.** Cross-backbone comparison on boundary reactions, five folds each, mean ± sd across folds. ΔAUROC is the per-fold change from the GNN-only score to the reconciled score. Gemma 4-31B-IT and Qwen3-32B share one GNN baseline and the other three share another because they were run on different fold draws. Backbones: Gemma 4-31B-IT [30], Qwen3-32B [31], Llama3-OpenBioLLM-8B [32], a biomedical fine-tune of Llama 3 [33], GPT-OSS-20B [34], DeepSeek-R1-Distill-Qwen-32B [35].

| Backbone | GNN-only | Reconciled | ΔAUROC |
| --- | --- | --- | --- |
| Gemma 4-31B-IT | 0.507 | 0.559 | $+0.0516 \pm 0.0311$ |
| Qwen3-32B | 0.507 | 0.521 | $+0.0137 \pm 0.0125$ |
| OpenBioLLM-8B | 0.504 | 0.504 | $-0.0001 \pm 0.0031$ |
| GPT-OSS-20B | 0.504 | 0.503 | $-0.0014 \pm 0.0193$ |
| DeepSeek-R1-32B | 0.504 | 0.473 | $-0.0317 \pm 0.0370$ |

### 4 Discussion

#### 4.1 Summary

We set out to test whether propagating patient transcriptomes across the metabolic network improves reaction activity prediction over scoring each reaction from its own genes. We cannot answer that question with the experiments we ran, and establishing why is the contribution.

Under expression-derived supervision the framework reached AUROC 0.9864 and 0.9839. Those numbers do not survive audit. Half the target is a threshold of the model’s own input, recoverable at 1.000 without any model; the other half is a random vector the model memorizes at 0.929 because patient-level splitting never hides the reaction axis. Outputs are near patient-invariant at 0.995, subsystem concordance is 3 of 8 and 0 of 5, and the model scores at chance on two external cohorts including a same-platform control.

Under input-independent supervision the framework never got the input. The released feature tensors for that cohort are uniformly zero, independently trained models agree on nothing patient-specific, and all three archived configurations sit at the no-signal null of the dispersion ratio while a model with verified features sits at more than twice it. The configuration comparison and the relational-edge gain built on those runs are withdrawn. Retrained on verified features, the model reaches 0.5800, below the 0.6342 that ranking by raw expression achieves with no model, and below the 0.6085 that an indicator carrying no expression information achieves.

Exactly one configuration in this study was run under conditions we can verify, and it did not exceed the raw-expression baseline. We state that plainly because the alternative, reporting the 0.6637 of the withdrawn three-edge run, would have been reporting a number produced without patient data.

#### 4.2 Recommendations

Four practices would have caught these failures, and all four are inexpensive.

#### Report the raw-expression baseline, and an information-free floor beside it

Both cost nothing and both are available on every cohort. The baseline is the minimum a scorer must beat to have shown that it adds anything to a univariate ranking of its own input. It does not by itself show that a model uses the network, for which a parameter-matched graph-free control is required and which we did not run. The indicator baseline shows how much of the target is prevalence structure rather than transcriptomic signal. On our cohort the indicator supplies 0.6085 of the 0.6342, and our verified configuration falls below both.

#### Use labels that are not a function of the model input, and audit their provenance

Where measured activity is unavailable, labels from an independent reconstruction are preferable to a threshold of the input matrix. But independence is not enough on its own: our own label vector contained 4,675 random entries through every experiment reported here without our noticing. Reproducing a stored label vector from its generating code should be a routine deposit check.

#### Test whether the model uses its input

Cross-validation, multi-seed replication, early stopping and tight standard deviations were all present in our archived runs and all reported normal behavior on models that were ignoring their inputs entirely. The cheapest sufficient check is cross-model agreement on patient deviations: train twice, center per reaction, correlate the residuals. It requires no ground truth and no uncertainty machinery. Where MC-Dropout is already in use, the dispersion ratio adds a second reading for the cost of a division, provided it is compared against its simulated null rather than against 1. We also suggest reporting the median inter-patient output correlation, with the caveat that in our hands it flagged both failures and separated neither.

#### Evaluate across consortia before claiming generalization

A same-platform cohort from a different consortium is the cheapest way to separate platform effects from everything else, and in our case it was decisive.

### 4.3 Limitations

The largest limitation is that this study contains no positive result about metabolic modeling. What we can support is a set of measurements about evaluation. Readers looking for evidence that graph structure helps reaction scoring will not find it here, in either direction.

Supervision is the second limitation, and we treat it as a finding rather than a caveat. For BRCA and LUAD the labels are partly a threshold of the model input and partly random. For the CRC cohort the labels are independent of the input, but they are derived from 11 NCI-60 cancer cell-line reconstructions, only one of them colorectal, with presence in a model treated as activity and 2,140 to 2,690 reactions matching per model. They are a coarse, tissue-mismatched proxy, and a scorer could be penalized for being right about colorectal biology. They remain the best input-independent target we could construct, and the raw-expression baseline is computed against the same labels, so the comparison between model and baseline is fair even where the labels themselves are poor. No cohort here is evaluated against measured reaction fluxes; establishing that any scorer in this family recovers metabolism requires validation against ^13^C metabolic flux analysis or targeted metabolomics, and that is the necessary next step.

Our verified retraining is at reduced scale: 100 of 624 patients, three folds, hidden dimension 64, on CPU. It is consistent across three variants but it is not the full-capacity experiment, and we do not claim it settles what a full-capacity model would do. The three-channel feature set used by the withdrawn run was never archived and cannot be recovered.

We ran no parameter-matched graph-free control, and we identify that as the single most informative missing experiment in this study. Without it, beating or failing to beat a univariate expression ranking says nothing about whether the network contributed, because a model with no edges at all would also depart from a univariate ranking. Any future claim that relational structure helps should be made against a graph-free model of matched capacity, not against the input baseline alone.

We also did not compute a dispersion ratio for the verified retraining, which would have been the matched positive control that Table 3 lacks; its per-patient predictions were not retained. The control we do report is from a different cohort with a different label rule and a different architecture, and it establishes only that the statistic can rise above its null.

We report no calibration metric; MC-Dropout outputs are uncertainty-aware, not calibrated. The empirical-Bayes discriminator used a simplified adjustment rather than the full estimator and constrains rather than excludes its mechanism. The proteomic channel is zero-filled throughout. The LLM module rests on five backbones evaluated on scores that were not patient-specific. The dispersion ratio assumes MC-Dropout samples are the only stochastic component at inference and that the *T* passes are independent, and it is normalized by each model’s own dropout magnitude, so it is not comparable across models with very different *σ*. It is a screening diagnostic, not a proof of input dependence: a value at the null warrants investigation rather than rejection, and in this paper the conclusion rests on the feature files and on cross-model agreement, with *ρ* as corroboration.

### 4.4 Relationship to other work

scFEA [37] estimates cell-wise fluxes on a 169-module reduced map rather than the full network, targets continuous flux rather than binary activity, and operates on single-cell data. FlowGAT [38] predicts essentiality on a 444-node mass-flow graph derived from *E. coli* iML1515, and its mammalian successor [39] extends flux-sampling graphs to Chinese hamster ovary and mouse models, again targeting gene essentiality rather than reaction activity. CHESHIRE [40] predicts missing reactions purely from network topology; CLOSEgaps [41] adds molecular-structure similarity and Multi-HGNN [42] is multi-modal, but none of the three uses patient omics and all three target gap-filling. HeteroGATomics [43] applies heterogeneous attention to patient-level classification rather than to a metabolic network. DeepMeta [44] predicts metabolic gene dependency with graph attention over an enzyme–gene network. TROPPO [45] wraps existing LP-based scorers rather than replacing them. Pan-cancer characterization of the metabolic reaction network [46] reconstructs models for over 4,000 tumors and establishes the biological motivation for patient-level scoring, though its per-sample reaction sets are themselves expression-derived and so cannot serve as an independent target. To our knowledge no published method combines patient-level expression input with reaction-level output on the full human reference network, though this is a claim about the literature we surveyed rather than one we can establish positively.

## 5 Conclusion

We asked whether propagating patient transcriptomes across the Recon3D graph improves reaction activity scoring over per-reaction independence. The honest answer from this study is that we do not know, and that the experiments we believed had answered it had not.

When activity labels are derived by thresholding the same expression matrix supplied to the model, apparent accuracy approaches the ceiling and carries no biological information. The input alone reproduces half the target, the other half is random and is memorized, outputs barely vary between patients, subsystem rankings largely fail to recover known phenotypes, and the model transfers to no external cohort, including one on the same platform. When labels are drawn instead from an independent reconstruction, the archived models turn out to have received no patient input at all. The one configuration we retrained on verified features falls below a baseline that uses no model and below an indicator that uses no data.

What we offer, then, is a documented case and the checks it taught us to run, not a method. Compare against the raw input feature, and against an indicator that carries no information at all, so that you know what your target is worth. Report how much the output varies between patients. Verify that the output depends on the input, by training twice and asking whether the two models agree about any patient. Replay your stored label vector from the code that generated it. Each costs minutes; none of them was part of our protocol; and between them they account for every result in this paper that we have had to withdraw. We release them with the curated cohort so that any method in this area, ours emphatically included, can be held to them.

Whether relational structure in the metabolic network helps patient-specific reaction scoring remains open. Answering it requires a model that demonstrably reads its input, evaluated against a parameter-matched graph-free control, on labels whose provenance has been replayed. None of those three conditions held for any experiment reported here, which is why we make no claim about it.

## 6 Software and Data Availability

Source code is available at https://github.com/thiptanawat/MetaGNN-Framework under the MIT License and archived at https://doi.org/10.5281/zenodo.21217569. Version 2 of the code record, which accompanies this manuscript, adds a diagnostics/directory containing the five checks of Section 2.7 as standalone scripts (raw-expression and indicator baselines, inter-patient correlation, cross-model agreement, dispersion ratio, and the label replay that reproduces the randomized partition of Section 3.2). Each prints the corresponding number in this paper when pointed at the deposited artifacts. Version 1 of the record predates this audit and does not contain them.

Processed inputs for the TCGA-CRC 624-patient cohort, including the rebuilt GPR-mapped reaction features described in Section 3.4, the HMA-derived labels and the fixed cross-validation splits, are deposited on Zenodo under CC-BY-4.0 at https://doi.org/10.5281/zenodo.21217579, together with the per-patient BRCA score archive and label vector used for Table 2, Table 3 and Figures 4 and 5. Readers should use version 2 of that record, which accompanies this manuscript. Version 1 (July 2026) contains a superseded 220-patient cohort that we have since found to be synthetic and the all-zero feature tensors described in Section 3.3; it remains accessible for the record but should not be used. The three-channel feature set used by the withdrawn experiment of Section 3.3 was not archived and cannot be released.

METABRIC and CPTAC-BRCA are not redistributable under their respective data-use terms. Download wrappers that reproduce our exact preprocessing from the original sources are included in the code repository.

## Declarations

### CRediT Author Contribution Statement

**Thiptanawat Phongwattana**: Conceptualization, Methodology, Software, Formal analysis, Investigation, Data curation, Writing – original draft, Writing – review & editing, Visualization. **Jonathan H. Chan**: Conceptualization, Methodology, Validation, Resources, Writing – review & editing, Supervision, Project administration, Funding acquisition.

### Ethics and Data Use

This study used de-identified, publicly available human genomic data obtained from The Cancer Genome Atlas via the NCI Genomic Data Commons, from cBioPortal, and from the Clinical Proteomic Tumor Analysis Consortium, in accordance with the data-use policies of each resource. No new human or animal data were collected. Institutional review board approval was not required.

## Acknowledgments

We are especially grateful to the IC2 Laboratory, School of Information Technology (SIT), King Mongkut’s University of Technology Thonburi (KMUTT), for experimental and supplementary tool support. We thank the TCGA Research Network, the Genomic Data Commons, the CPTAC consortium, the Human Metabolic Atlas and the Virtual Metabolic Human project for making the underlying resources publicly available.

